## Supplementary figures and images for "Exploring causal effects of smoking and alcohol related lifestyle factors on self-report tiredness: a Mendelian randomization study"

### Fig A.tif

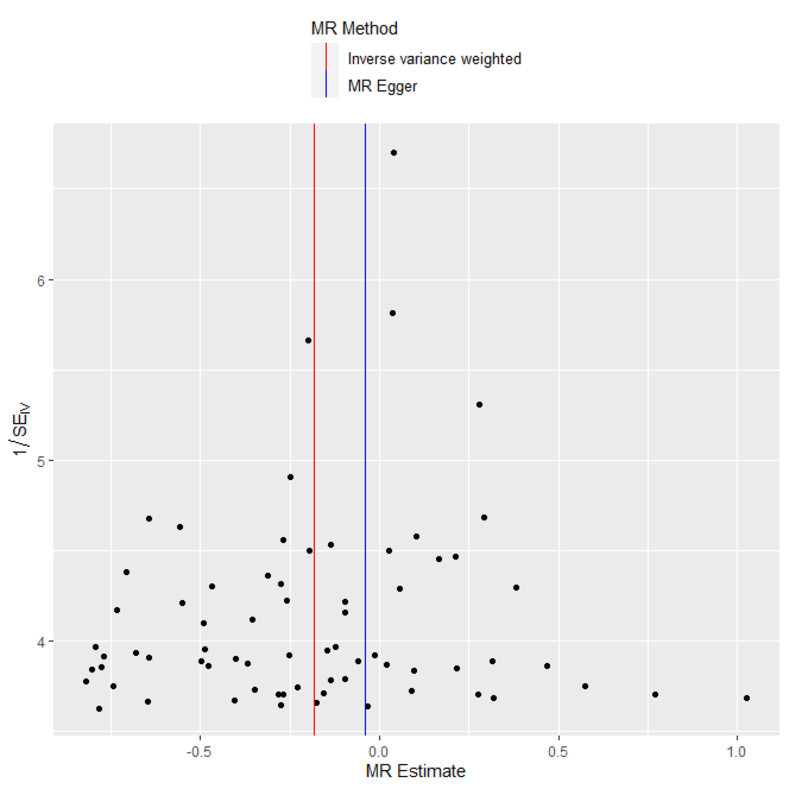

### Fig B.tif

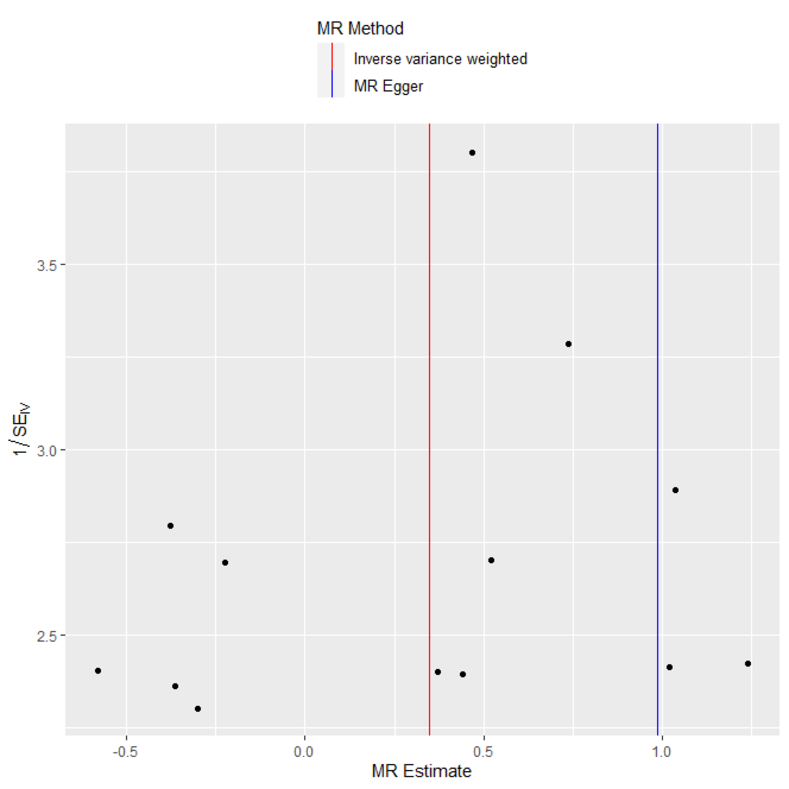

### Fig C.tif

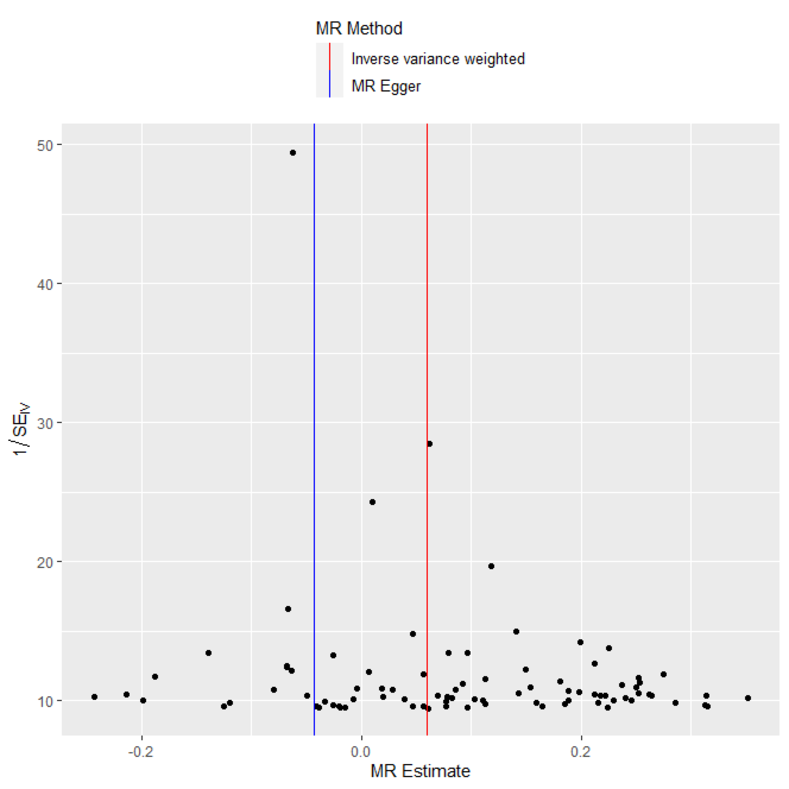
